# Improvement of eukaryotic proteins prediction from soil metagenomes

**DOI:** 10.1101/2021.11.10.468086

**Authors:** Carole Belliardo, Georgios D. Koutsovoulos, Corinne Rancurel, Mathilde Clément, Justine Lipuma, Marc Bailly-Bechet, Etienne G.J. Danchin

## Abstract

**Background:** During the last decades, shotgun metagenomics and metabarcoding have highlighted the diversity of microorganisms from environmental or host-associated samples. Most assembled metagenome public repositories use annotation pipelines tailored for prokaryotes regardless of the taxonomic origin of contigs and metagenome-assembled genomes (MAGs). Consequently, eukaryotic contigs and MAGs, with intrinsically different gene features, are not optimally annotated, resulting in an incorrect representation of the eukaryotic component of biodiversity, despite their biological relevance.

**Results:** Using an automated analysis pipeline, we have filtered 7.9 billion of contigs from 6,873 soil metagenomes in the IMG/M database of the Joint Genome Institute to identify eukaryotic contigs. We have re-annotated genes using eukaryote-tailored methods, yielding 8 million eukaryotic proteins. Of these, 5.6 million could be traced back to non-chimeric higher confidence eukaryotic contigs. Our pipeline improves eukaryotic proteins completeness, contiguity and quality. Moreover, the better quality of eukaryotic proteins combined with a more comprehensive assignment method improves the taxonomic annotation as well.

**Conclusions:** Using public soil metagenomic data, we provide a dataset of eukaryotic soil proteins with improved completeness and quality as well as a more reliable taxonomic annotation. This unique resource is of interest for any scientist aiming at studying the composition, biological functions and gene flux in soil communities involving eukaryotes.

**Key Points**

- Eukaryotic micro-organisms have key roles in soil microbial communities and ecosystems.
- Eukaryotic genes and proteins are incorrectly represented in metagenomic libraries due to inappropriate methods dedicated to prokaryotes.
- We have improved the quality and completeness of the *de novo* proteins prediction and taxonomic annotation of eukaryotic organisms from soil or plant-associated microbiome.

## Background

Soil-dwelling microorganisms play essential roles in biological functions related to human and Earth health in both managed and natural ecosystems [1]. They are engaged in countless processes, from chemical engineering to environmental change, or nutrient cycling through soil fertilisation or carbon sequestration [2]. In recent years, the rise of metagenomics has expanded our understanding of the genetic diversity of microorganisms in many different complex environments, including soil and plant-associated microbiomes [3]. First, metabarcoding has highlighted the high variability of microbial communities across different soil samples [4]. Then, shotgun metagenomic sequencing has allowed the discovery of unknown microorganisms [5]. Recently, several efforts have focused on the *de novo* assembly of bulk metagenomic sequencing reads into metagenome-assembled genomes (MAGs) or contigs, uncovering these microbe’s genetic content and augmenting our understanding of their metabolic pathways [6, 7, 8].

The soil is the most complex microbiome due to the extremely high diversity of organisms, of their complex inter-kingdom interactions and high range of environmental conditions observed between samples; in contrast, the human gut microbiome is more homogeneous among individuals due to constant physiological conditions. Indeed, soil can differ in abiotic, as pH, moisture and texture, and biotic properties, as fauna and flora [9, 10], even if samples are a few millimetres apart [5]. Therefore, the soil microbiome is rather a patchwork of distinct microenvironments, containing many microbial guilds with disparate metabolic abilities living at different locations and depths [11].

The number of individual taxa in one gram of soil is estimated at several thousand on average and cover all different superkingdoms of life (Bacteria, Arachaea and Eukarya). Most soil metagenomic studies are focused on bacteria that constitute a prominent part of the microbiome in individual numbers, even though eukaryotes often account for a comparable biomass [3]. The composition and diversity of eukaryotic microorganisms in soils are expected to be higher than, and different, from other ecosystems but are still mostly unknown [9, 10, 2].

However, soil eukaryotic microorganisms fulfill essential functions in ecosystems. They participate in the biochemical balance, such as the carbon sequestration in the soil, and thus could be relevant to modeling environmental issues [12]. Eukaryotic microorganisms living in soils can display all trophic types and lifestyles, and affect the biodiversity and health of macro-organisms constituting the soil fauna and the flora. Some species play a role in regulating the population and shaping other communities composition by predation, as in the case of phagotrophic eukaryotic microbes, which also play a key role in soil fertilisation [13]. Some other eukaryotes are pathogens of plants and animals (including livestock or even humans): they are present as propagules in soil, acting like reservoirs, and can cause tremendous health or economic damage [14].

A substantial part of these soil microbes is involved in nutrient cycling as decomposers or symbionts associated with plants. One of the most outstanding cases is mycorrhizal fungi which live symbiotically with 90% of the vascular plants on Earth [15]. The mutualistic interaction with these eukaryotic microorganisms from the rhizosphere provides nutritive and protective benefits for plants, giving them a strong agronomic interest [16, 17]. These fungi are also engaged in other ecological services with high environmental interest such as reforestation [18, 19]. Even though these microorganisms are useful in plant-related management, their genetics and ecology are still poorly understood. Soil metagenomics could open a way to address these issues.

Despite their importance, soil eukaryotes are neglected and not well represented in public metagenomic data. The largest publicly available resource for soil (and other) metagenomes is the Integrated Microbial Genomes & Microbes (IMG/M) database of the Joint Genome Institute (JGI) [20]. In this resource, standard pipelines are used to assemble and annotate contigs and genomes from environmental metagenomic shotgun reads. One major limitation concerns the eukaryotic component of these soil metagenomes. Indeed, the gene prediction tool used by default for all contigs assembled from metagenomes is Prodigal [21], a software tailored for prokaryotes. However, gene structures and features are different in eukaryotes and using prokaryotic tools to predict eukaryotic genes can lead to incomplete, erroneous and discontinuous gene sequences, and hence proteins [22]: a trivial example is that no intron can be predicted by Prodigal. Furthermore, in the JGI standard procedure, the taxonomic annotation assigned to proteins is based on the single best hit from a homology search against the NCBI’s nr library, which may itself be taxonomically mis-annotated, or result in annotation at the species or strain level despite of metagenomics sequences having low identity with their nr homologs.

These procedures make sense given the volume of metagenomic data re-treated by IMG/M, but, as a consequence, eukaryotic proteins are neglected in these soil microbiome data, and may truncated and assigned an unreliable taxonomic annotation. These suboptimal sequences and taxonomic annotations then negatively impact any research on the eukaryotic component of the soil. To circumvent this problem, we have first detected eukaryotic contigs in soil microbiomes using a k-mer based approach. On the identified eukaryotic contigs, we then re-predicted genes and proteins using annotation methods tailored for eukaryotes. We show that the newly predicted proteins are longer and constitute a more comprehensive representation of the pool of eukaryotic proteins in the soil. Using these improved protein predictions, we re-assigned taxonomic information based on a last common ancestor (LCA) approach from homology search against the NCBI’s nr. This new dataset improves protein sequence quality and completeness, as well as the reliability of the taxonomic information, and represents a unique resource to decipher and study the pool of proteins eukaryotic present in the soil.

## Data description

### Methods

#### Data collection

We used publicly available assembled metagenomic data from shotgun sequencing reads of the IMG/M database of the JGI [20]. We collected metagenomes including 5,989 ‘Terrestrial’ samples in the environmental metagenomes category and 884 plant-associated metagenomes in the host-associated category (available data 2020, October see supp. data 1). The data acquisition was performed via the IMG/MER Cart genome portal. For each metagenome, the JGI provides a set of files from pre-computed analyses that are useful to sort, filter and describe data. Because we anticipated substantial differences in the relative proportions of eukaryotic species present in the terrestrial and the host-associated categories, these two datasets were processed separately to minimize potential biases. For a more convenient processing of the huge amount of data, the metagenomes from terrestrial samples were splitted in two batches; ‘Terrestrial 1’ contains 3,601 environmental metagenomes added between december 2009 and january 2019 and the second one called ‘Terrestrial 2’ contains 2,388 data added between February 2019 and August 2020.

#### Data curation and quality control

Starting from assembled contigs, we combined all genomic fasta files by datasets and obtained 6 and 1.9 billion contigs from terrestrial and plant-associated categories, respectively. The quality of assembled contigs is highly heterogeneous between metagenomes due to variation in sequencing technologies, experimental protocols, pipeline version used and biological features. Probably because most data initially consist of short sequencing reads, half of the contigs were shorter than 296bp (Table 1). These short contigs increase the volume to be processed and are unlikely to contain complete genes, hence providing no more information on gene diversity [23]. Thus, we filtered data on assembly quality, and kept contigs at least 1kb long or containing at least three genes. Only 763 million contigs (10%) passed this filter and were retained for further analysis. These remaining contigs are distributed in 6,611 metagenomes: hence data from 262 starting metagenomes were entirely removed due to a too high level of fragmentation. This quality filtering drastically reduces the dataset volume and ensures we only work with contigs on which complete genes can be predicted.

**Table 1.**
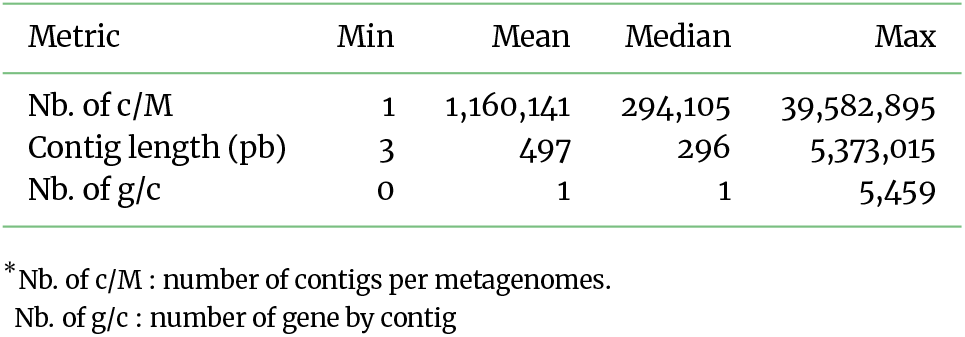
Metrics to assess the quality of the 6,873 ‘Terrestrial’ and ‘plant-associated’ metagenome-assembled genomes datasets from the IMG/M server of the JGI.

#### Detection of contigs from eukaryotic organisms

The JGI provides taxonomic information for genes but no information is provided for contigs. We then scanned all the contigs and identified those from eukaryotic origins using Kraken2, a taxonomic classification tool based on exact kmer matches to achieve as fast and accurate as possible processing of large data sets such as those from metagenomics analyses [24]. As a reference database, we combined all RefSeq libraries of complete genomes [Archaea, Bacteria, Plasmid, Viral, Human, Fungi, Plant, Protozoa, nt] [25], and ran Kraken2 with default parameters. This allowed assigning taxonomic information to 82% of contigs, among which 113 million were classified with a eukaryotic TaxID (18%).

#### Eukaryotic gene prediction

For all contigs identified as eukaryotic by Kraken2, we used Augustus (v3.3), a software dedicated to *de novo* eukaryotic gene prediction [26]. The gene structure is complex in eukaryotes and changes across species [27]. Thus, Augustus provides *ab initio* models for 73 different species (Fig. 1) and one must be selected to perform gene prediction. Due to the conservation of genomic features across closely related organisms, we assigned, for each eukaryotic contig, a model based on its Kraken2 taxonomic annotation. Note that this model selection step does not aim at a definitive taxonomic annotation; here we predict as best as we can putative eukaryotic genes that will then be filtered by a taxonomic annotation approach. For the selection of the phylogenetically closest model for each putative protein, we used a custom python script ([see code availability](https://github.com/CaroleBelliardo/EukaProt_in_PublicSoilMetag)) which functions as follows:

- First, we browsed the 73 model species tree from the leaves to the root assigning a non-ambiguous parental taxonomic term to each model species as long as no bifurcation with a branch leading to another model species was found (Fig. 1). For example, in the plants, *Arabidopsis thaliana* is the sole representative of the Brassicales; so the Brassicales parental term was associated with the *A. thaliana* model. Consequently, we used the *A. thaliana* Augustus model for all eukaryotic contigs assigned with a taxonomic ID belonging to the Brassicales branch. Similarly, *Homo sapiens* is the only representative of mammals, so any contig identified by Kraken2 as a mammalian organism will be assigned the *H. sapiens* model.

**Figure 1.**
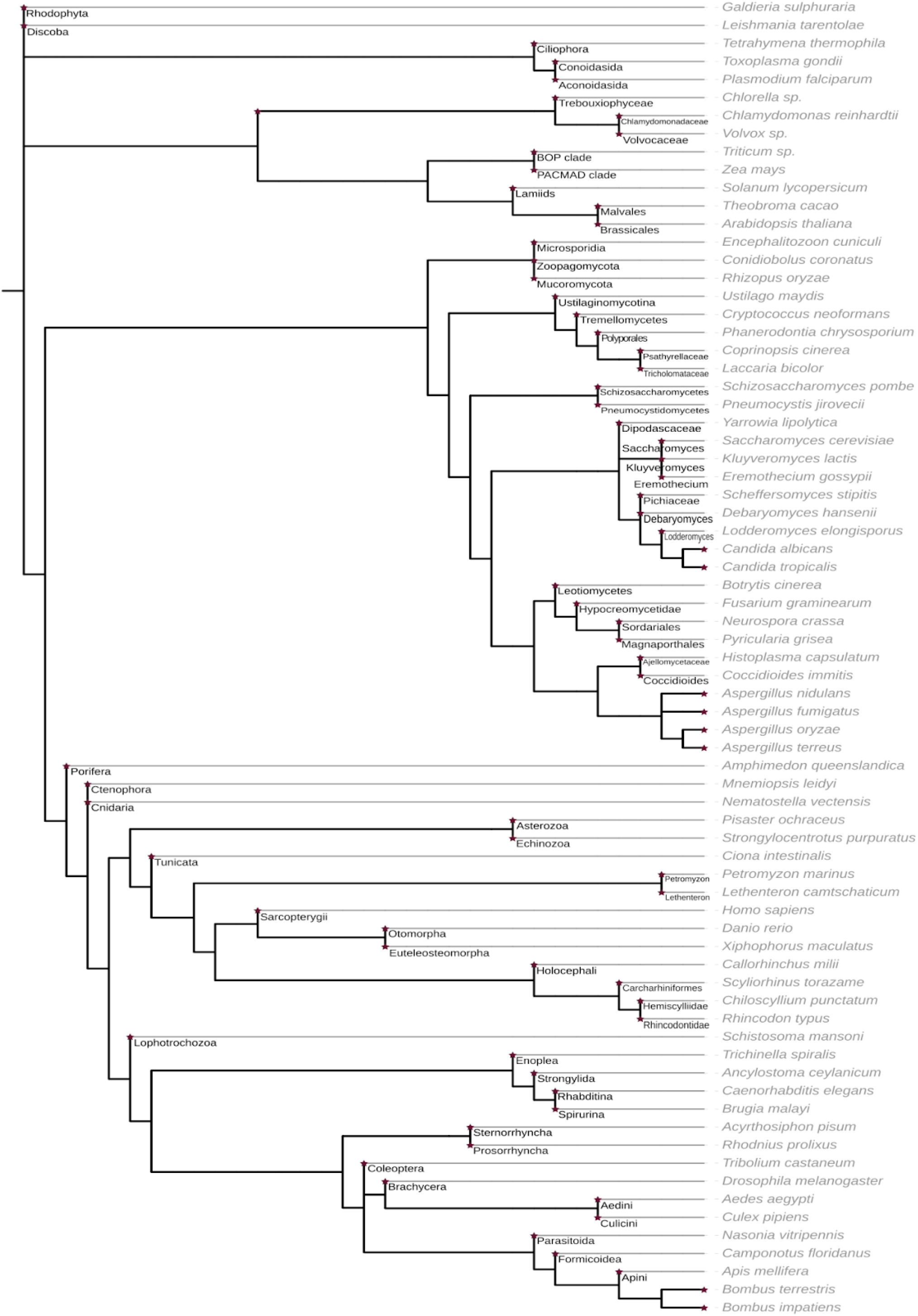
Phylogenetic tree of Augustus *ab initio* models showing, by a red star, the deeper taxonomic node used in the first step of the contig model selection.

At this point, an Augustus gene prediction model could be assigned to 7,1% (8 million) of contigs. The rest of the contigs (>100 million) could not be assigned an unambiguous closest model species because they belonged to a bifurcating branch in the tree leading to several equally close model species.

- Therefore, in a second step, for all these remaining eukaryotic contigs, we selected among the child branches the most frequently assigned model in the whole dataset the contig belongs to (i.e. Plant-associated, Terrestrial 1 or Terrestrial 2) at the previous step (first pass). To continue the previous example, the next more ancestral branch in the phylogeny of Brassicales is the clade ‘Malvids’ that displays a polytomy of eight child branches of which only two are bearing gene model species (Malvales and Brassicales). Hence, no model could be unambiguously assigned to contigs with a Malvids taxonomic ID other than Malvales or Brassicales. Therefore, all contigs from other Malvids orders are processed with the most frequently assigned species model for each of the three datasets (Fig. 2). For example, they are processed with the cocoa gene model (Malvales, *Theobroma cacao*) in the dataset Terrestrial 1, or the *Arabidopsis thaliana* gene model in the plant-associated dataset (Fig. 2). The distribution of contigs across the models is available in [supplementary data Fig. 1](https://data.inrae.fr/dataset.xhtml;jsessionid=79633dcca30d147adbb76f0c1c13?persistentId=doi%3A10.15454%2FE2VTRB&version=DRAFT).

**Figure 2.**
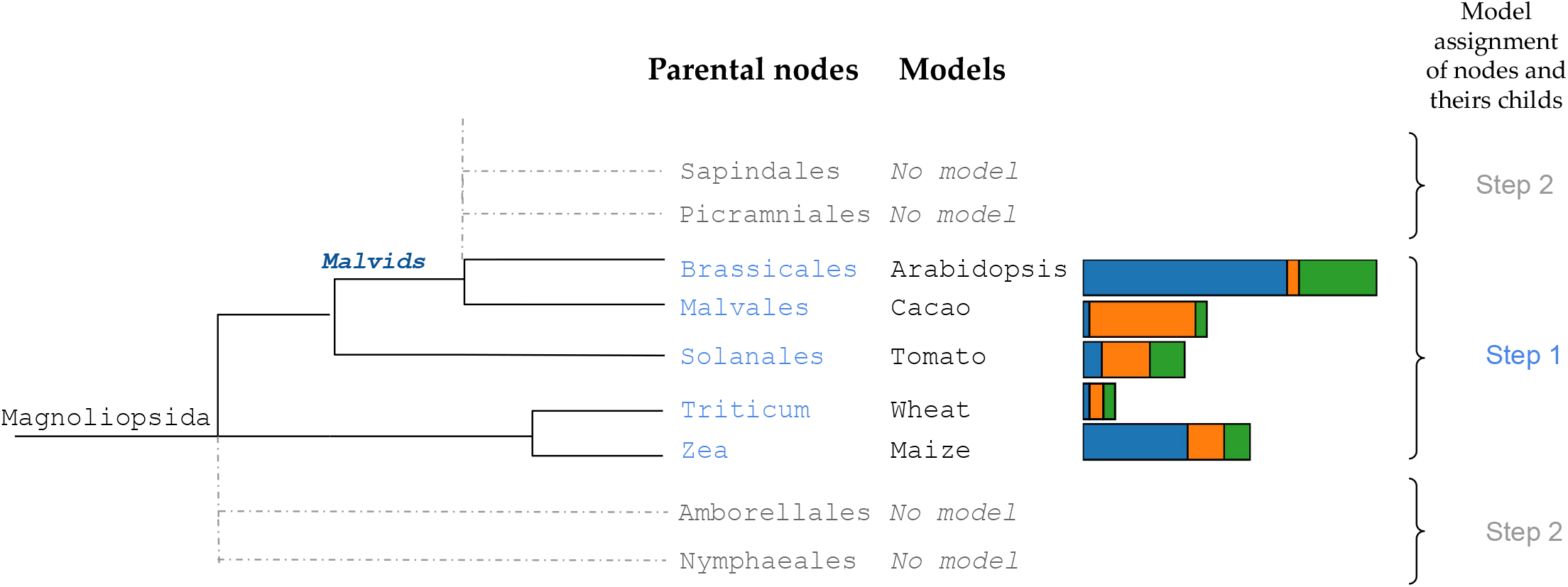
Phylogenetic tree focused on Magnoliopsida clades displaying the Augustus model distribution supporting the assignment of *ab initio* gene model by dataset (blue= Plant-associated, orange=Terrestrial 1, green=Terrestrial 2).

Overall, our pipeline allowed assigning an Augustus model to over 100 million possibly eukaryotic contigs. The most assigned models are metazoa and viridiplantae models, with respectively 49% and 44% of contigs in plant-associated metagenomes and 76% and 16% in terrestrial data. In both datasets, we assigned fungal models to 6.5% of contigs (see [suppl. table 1](https://data.inrae.fr/dataset.xhtml;jsessionid=796a90dd0dad84f5f6973c5a60a6?persistentId=doi%3A10.15454%2FE2VTRB&version=DRAFT)); and the majority of other contigs were assigned to SAR, Discoba or Rhodophyta models. Although these last taxonomic groups were assigned at a relatively low ratio, this corresponds to tens or hundreds of thousands of contigs. Unsurprisingly, the less assigned are gene models of aquatic animals such as some benthic animals, sharks, or also lamprey models. At this point, we could not assess whether the numerous assignments to the metazoan and plant models came from miss-annotated contigs or contamination, therefore further analyses were performed after protein prediction.

Then, once a model species has been assigned to contigs we run the eukaryotic gene predictor Augustus [26], with default parameters, which allows predicting 93 million protein-coding genes. The number of proteins predicted per contig is between 1 to 410 with 2 protein predicted per contig on average for all datasets together. Consistent with the contig model assignment across kingdoms, the highest numbers of proteins were predicted for contigs assigned to metazoan and viridiplantae Augustus models. Moreover, we predicted 8.7 million proteins with fungal models and 1.8 million with different protist models (Fig. 3 A, [suppl. table 2](https://data.inrae.fr/privateurl.xhtml?token=5fe3ff97-c716-41ff-98a0-f6e9aa0631d3)).

**Figure 3.**
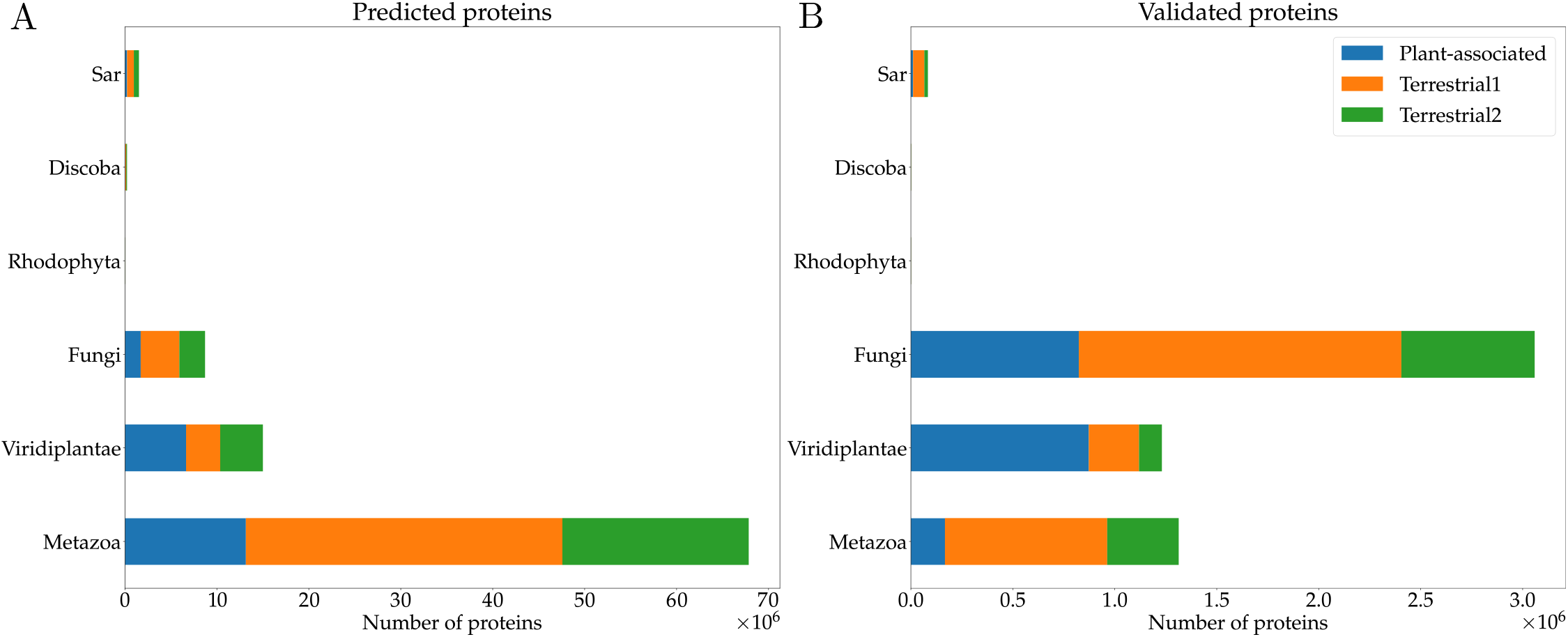
Number of Augustus-predicted proteins and their taxonomic distribution per Augustus model kingdom by dataset (A) on all contigs (B) on non-chimeric eukaryotic contigs validated by Diamond (blue= Plant-associated, orange=Terrestrial 1, green=Terrestrial 2).

#### Confirmation of eukaryotic origins and improvement of the taxonomic information

To exclude false-positive eukaryotic classification and assign a more reliable taxonomic annotation to the proteins than simple inheritance from the Kraken2-based contig annotation, we used Diamond with the LCA ‘last common ancestor’ mode for all predicted proteins [28]. The homology search was run with an E-value threshold of 10^−6^ using the NCBI nr library as a reference protein database. In the LCA mode, Diamond will assign an NCBI taxonomic identifier based on the last common ancestor of all the hits with a score not diverging by more than 10% from the best hit score. Using an LCA approach constitutes a substantial gain in taxonomic annotation reliability compared to approaches based on the best blast hit alone, this single best hit being potentially mis-annotated itself, or sharing low identity with the query sequence. Then, many “chimeric” contigs contained proteins from both eukaryotic and non-eukaryotic or untraceable / unknown origin predicted by the LCA approach. To be stringent, we retained only proteins from contigs in which at least half of the predicted proteins were assigned an eukaryotic taxonomic annotation. In most cases, the contigs deemed chimeric contained eukaryotic proteins not reaching the 50% threshold, but the non-eukaryotic proteins had no taxonomic annotation, either because they returned no hit in the homology search, or an ambiguous annotation (e.g. ‘unclassified’ or ‘other’ NCBI taxid): there were few cases of chimeric contigs with both eukaryotic and prokaryotic proteins. In the end, 6 percent of the predicted proteins, spanning 4,060 metagenomes in all datasets, passed these filters and were considered as present on validated eukaryotic contigs (see [suppl. table 3](https://data.inrae.fr/privateurl.xhtml?token=5fe3ff97-c716-41ff-98a0-f6e9aa0631d3)). Eukaryotic proteins from potential chimeric, non validated contigs, are available for further study in [suppl. data fasta file](https://data.inrae.fr/privateurl.xhtml?token=5fe3ff97-c716-41ff-98a0-f6e9aa0631d3) with [suppl. data taxonomic data](https://data.inrae.fr/privateurl.xhtml?token=5fe3ff97-c716-41ff-98a0-f6e9aa0631d3). As expected for soil metagenomes, the numbers of proteins validated as belonging to non-chimeric eukaryotic contigs are close to zero for some contigs with Kraken2-assigned taxa from aquatic organisms (e.g. *Mnemiopsis leidyi*) or mammals parasitic organisms (e.g. *Encephalitozoon cuniculi*). The proportion of proteins predicted by Augustus model on non-chimeric eukaryotic contigs ranged between 0.6% and 5.7% for models of all kingdoms except Fungi, where this proportion reached respectively more than 33% and 50% for plant-associated and terrestrial datasets (Fig. 3 B, [suppl. table 4 & 5](https://data.inrae.fr/dataset.xhtml;jsessionid=799a51b1a004dabf1b7bd46fae55?persistentId=doi%3A10.15454%2FE2VTRB&version=DRAFT)). Therefore, a high proportion of proteins predicted with Metazoan Augustus models due to initial Kraken2 taxonomic annotation were eliminated by this Diamond-based filtering of non-chimeric eukaryotic contigs, and consequently, proteins from fungal models became the dominant part of validated proteins. An explanation for this important change in model proportion between the two steps of our analysis may be the following. A substantial number of contigs were probably assigned a eukaryotic taxonomy by Kraken2 based on a low number of k-mer matching with the eukaryotic target. The proteins predicted on these contigs were not assigned a eukaryotic annotation by Diamond but either a prokaryotic or unknown taxonomy. Applying a confidence score threshold to Kraken2 taxonomic predictions might have resolved part of these false positives but at the risk of augmenting the rate of false negatives, and thus missing some eukaryotic contigs. Therefore, our strategy was to first apply a highly sensitive Kraken2-based filter to retrieve as many putative eukaryotic contigs as possible and then to apply a very stringent Diamond-based filter to only retain non-chimeric contigs.

## Data Validation and quality control

To determine whether using Augustus in our pipeline allowed improving the eukaryotic protein predictions, we compared them to the proteomes obtained by the JGI for the same set of contigs. In non-chimeric, validated eukaryotic contigs, a total of 16 million proteins were predicted by Prodigal in 3,294,767 contigs from 3,980 metagenomes – vs. 5.6 million of proteins with our pipeline (see [suppl. table 6](https://data.inrae.fr/privateurl.xhtml?token=5fe3ff97-c716-41ff-98a0-f6e9aa0631d3)). However, the raw number of proteins can be misleading because while Prodigal predicted 2,5 billion amino acids, we predicted 1,9 billion amino acids in total with Augustus, suggesting although more proteins were predicted by Prodigal, they were much shorter. Augustus allowed predicting introns in 1,627,033 genes from 1,074,415 contigs; these intronic sequences span on average 17% of the gene length. In comparison, Prodigal is not able to predict introns and end its prediction when the first stop codon is encountered. Therefore, at least 28% (1.6/5.6 millions) of the proteins predicted by Augustus were necessarily incorrectly predicted by Prodigal, initially. Moreover with the high frequency of stop codons in the intronic regions due to less selective pressure on these genomic regions, most intron containing proteins are expected to be truncated by Prodigal. To further compare predictions from both tools, we used two metrics: *(i)* protein length distribution, and *(ii)* the recovery of universally conserved unique copy eukaryotic genes.

### (i) The length distribution of protein

First, we calculated and compared the distribution of protein lengths from Augustus vs. Prodigal. Proteins from Augustus are significantly longer than protein *de novo* predicted with Prodigal (Fig. 4; unpaired t-test, n=5.3 10^6^/n=9 10^6^ proteins, stat=1.994 10^3^, *p ≤* 10^−4^). These observations coupled with the higher number of proteins predicted by Prodigal, suggests that Augustus was able to predict introns and join together multiple exons to form more complete genes where Prodigal predicted multiple truncated genes : the average size of genes (in ext. proteins) in eukaryotes is larger than in prokaryotes, due to the evolution of genome complexity [29]. Furthermore, the length distribution is closer to a normal one with Augustus predictions than with Prodigal ones (Fig. 3 B), indicating a better quality of our new predictions. In this regard, Nevers et al., (2020) reports that a non-normal distribution of proteins length, as observed for these Prodigal predictions in eukaryotic contigs, is indicative of more truncated proteins caused by fragmented genomes and incorrect protein prediction[30].

**Figure 4.**
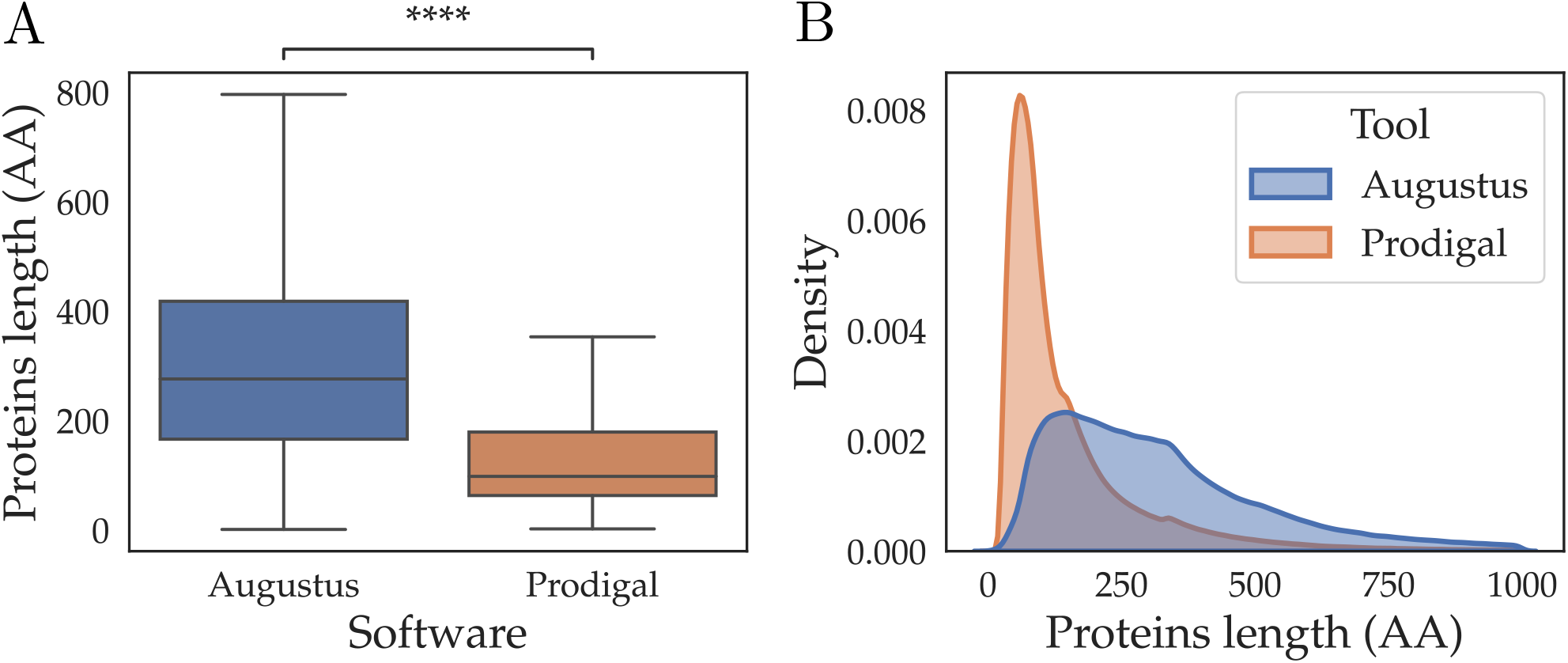
Distribution of protein lengths of Augustus prediction in blue versus Prodigal prediction in orange. Proteins from Augutus are significantly longer than those from Prodigal (see text).

### (ii) The recovery of universally conserved unique copy eukaryotic genes

To assess the improvement of our *de novo* protein prediction from eukaryotic soil microorganisms, we compared the proportions of near-universal single-copy orthologs retrieved for each metagenome with those provided by Prodigal in the same contigs using BUSCO (v.4.0.2) in protein mode with ‘*eukaryota_odb10*’ lineage [31]. Universally-conserved eukaryotic BUSCO proteins were identified in 1,093 metagenomes from the 4,060 metagenomes containing predicted eukaryotic proteins. This observation is not particularly surprising since *(i)* there are only 255 universally-conserved eukaryotic BUSCO genes, *(ii)* eukaryotes represent a minority of species in the soil and *(iii)* most eukaryotic genomes are only partially assembled from short-read bases shotgun metagenomic data.

The proportion of BUSCO genes found in complete length in metagenomes was significantly higher for the Augustus predictions than for the initial Prodigal predictions (Fig. 5; paired Wilcoxon-test, n=1,093 metagenomes, Complete stat=1.132 10^5^, *p ≤* 10^−4^; Fragmented stat=2.039 10^4^, *p ≤* 10^−4^). Similarly, the proportion of fragmented and missing BUSCO genes were significantly lower in Augustus predictions as compared to Prodigal predictions; this trend is identical for all datasets (see [supp. data, fig. 2](https://data.inrae.fr/privateurl.xhtml?token=5fe3ff97-c716-41ff-98a0-f6e9aa0631d3)). The BUSCO completeness scores from our Augustus gene predictions are as good or better than Prodigal for more than 98% of metagenomes. Furthermore, we have predicted more universal single-copy genes than Prodigal for 574 metagenomes, or more than half of the 1,093 metagenomes containing at least one BUSCO gene in one of both predictions. We observe an average improvement of 11.9% in the BUSCO completeness score, and genes are less fragmented in 510 metagenomes with an average of 8.5% lower proportion of fragments (see [suppl. table 7](https://data.inrae.fr/privateurl.xhtml?token=5fe3ff97-c716-41ff-98a0-f6e9aa0631d3)). The scores provided by BUSCO for these 1,093 metagenomes show a significant improvement of protein recovery and completeness for proteins from Augustus compared to those from Prodigal, indicating our pipeline has improved the quality of eukaryotic gene models in soil metagenomes.

**Figure 5.**
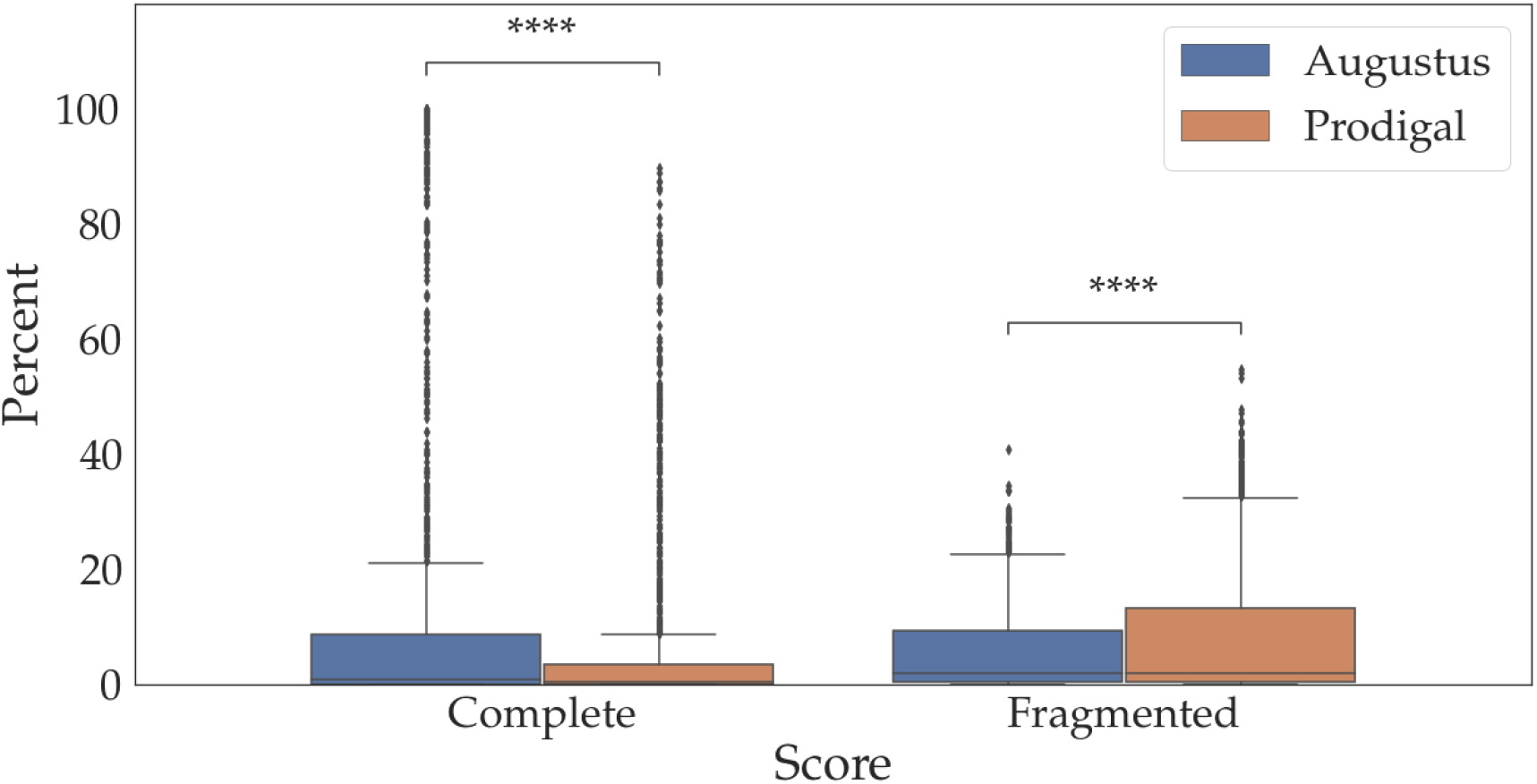
The Complete and Fragmented Busco score of of the 1,093 metagenome with single-copy universally conserved genes report a significantly better recovery of genes from eukaryotic microorganisms with Augutus than Prodigal (see text).

### Taxonomic assignment of eukaryotic proteins

Aiming at assigning a reliable taxonomic annotation to the proteins predicted by our pipeline, we used the last common ancestor algorithm of Diamond [28] and a recent (January 2020) NCBI nr database version [25]. In our approach, a taxonomic annotation is assigned based on the last common ancestor of all the hits whose scores did not diverge by more than 10% from the score of the best hit. This strategy contrasts with the default automated JGI procedure which assigns a taxonomic annotation based on the best blast hit only, regardless of the % identity. As a matter of fact, the last common ancestor approach is usually used to taxonomically assign sequences far related to the known organisms, such as ancient or actual metagenomic data [32, 24, 33, 34, 35]. Consequently, the improved quality and completeness of protein sequences combined with a more accurate taxonomic assignment method is expected to yield a more reliable taxonomic annotation.

This process allowed assigning a taxonomic annotation to 94% of the proteins predicted in non-chimeric eukaryotic contigs (Fig. 6). Most of them are assigned as fungal (e.i. 63%). These results are consistent with eukaryotic taxonomic distribution previously described in the literature, reporting Fungi as the most abundant eukaryotic microorganisms in studied soil [3, 36]. Actually, in soil metagenomes, fungal organisms are often second to bacteria in number and account for a comparable proportion of the biomass. The second most represented kingdom in our taxonomic assignment is Viridiplantae with 20% of annotated proteins. Not surprisingly this relatively high proportion is due to the plant-associated metagenomes, where nearly half of proteins are assigned as plant proteins (see 4,5,6). The following most represented is the SAR phylum (2%); this small percentage still represents several million proteins. In our taxonomic annotation, some proteins are still annotated as bacterial or viral during the final step and need to be carefully considered. Although we only retained contigs with at least 50% of eukaryotic genes, some misassembled chimeric contigs can still remain in the dataset. However,on these contigs only 0.01% of the predicted proteins were annotated with prokaryotic or viral lineages. Alternatively, because these viral and prokaryotic genes could also represent horizontal gene transfers within eukaryotic contigs, these genes were kept in the dataset. Without a major change in the relative representation at the kingdom or deep phylum level, this work enriched the taxonomic information provided for some eukaryotic microorganisms families by expanding the number of taxonomically-annotated proteins but also the diversity of taxonomic identifiers assigned. For example, we know that Arbuscular Mycorrhizal Fungi (AMF) are ubiquitous members of soil microbiota, and more particularly of the rhizosphere [15]. These eukaryotic microorganisms are plant symbionts with high impacts in several fields, mainly in agronomy due to a biostimulant and a bioprotection effect [37, 16], but they are also used to help in environmental issues such as cleaning-up of polluted soils or facilitating reforestation. As a matter of fact, in the original JGI data, only 8,065 AMF proteins were predicted in 6,048 contigs from 327 metagenomes with solely one species/TaxID assigned (i.e. *Rhizophagus irregularis*). Using our eukaryotic-centred gene prediction and taxonomic annotation pipeline, we expand the identification of AMF to 50,999 proteins in 48,726 contigs from 1,102 metagenomes. Moreover, this new annotation now covers 26 different taxa (from class to species) better representing the AMF diversity present in these soils (Fig. 7). The case of these pervasive eukaryotic microorganisms in the soil highlights the benefits of this work to improve the representation of eukaryotic organisms in public soil metagenomes.

**Figure 6.**
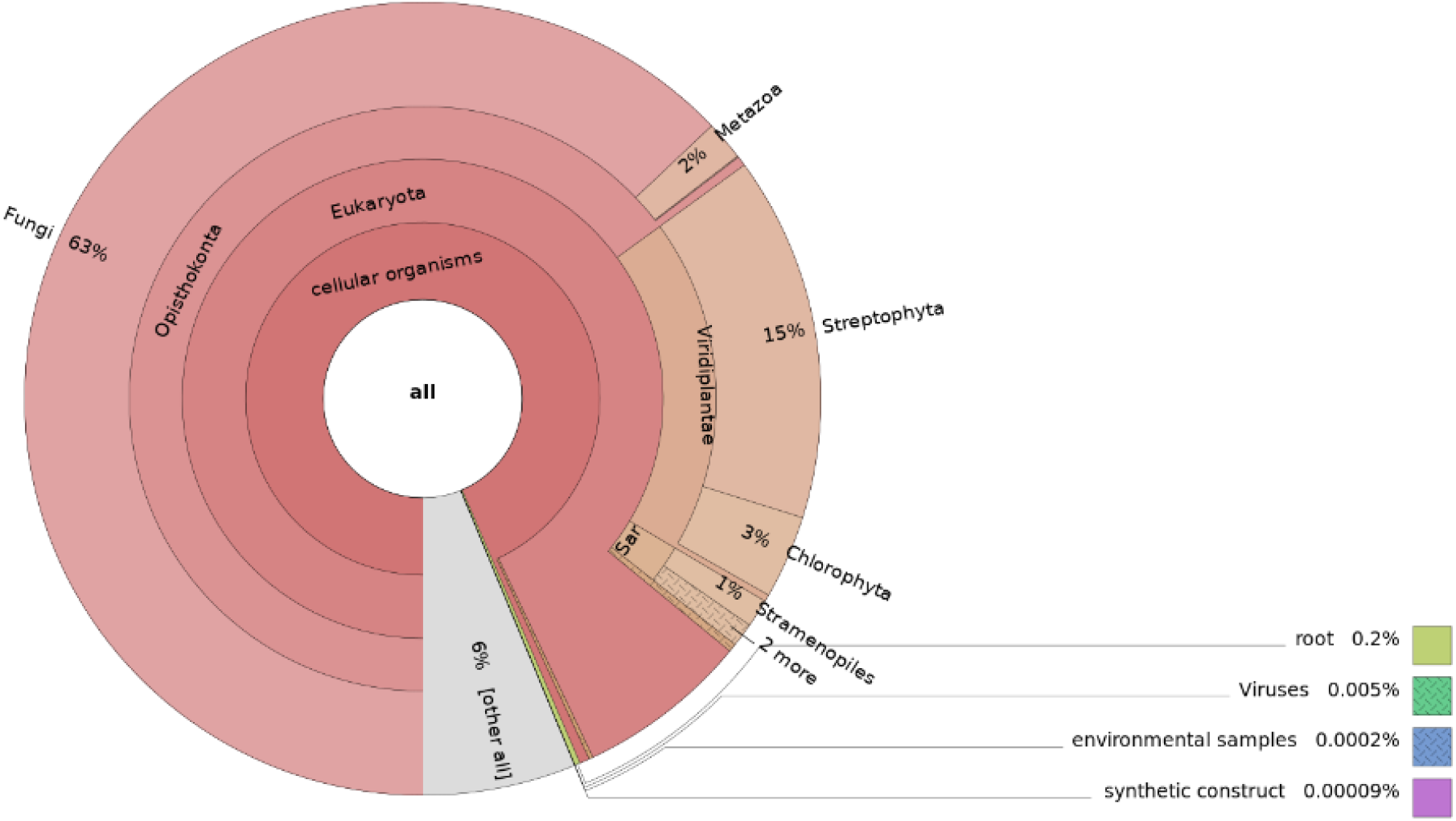
Krona representation of taxonomic assignment provided by the last common ancestor algorithm of DIAMOND for all the 5.6 million new eukaryotic proteins predicted by our homemade pipeline using Augustus (html file : available on [supp. data fig 3](https://data.inrae.fr/privateurl.xhtml?token=5fe3ff97-c716-41ff-98a0-f6e9aa0631d3)).

**Figure 7.**
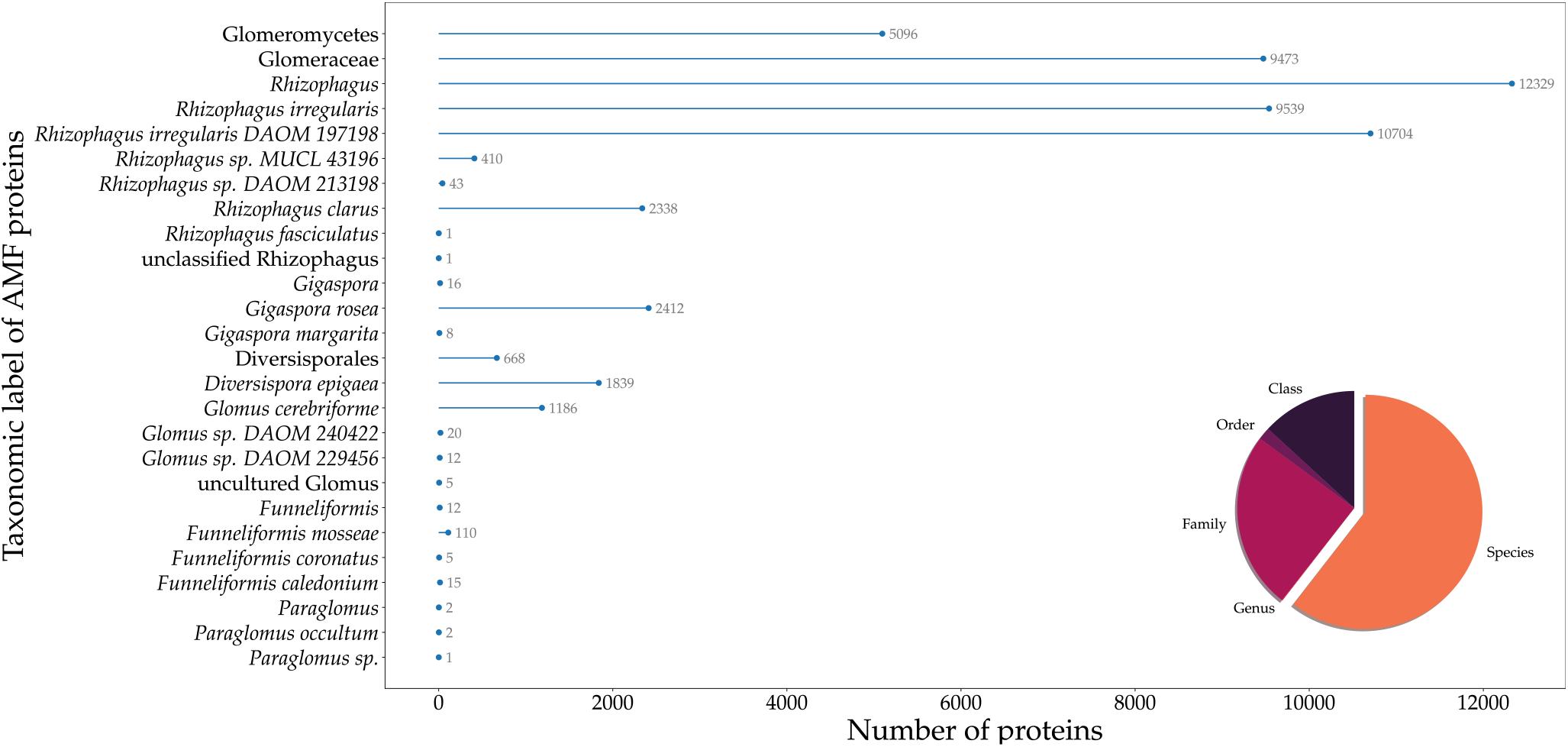
Annotated taxa of Arbuscular Mycorrhizal Fungal proteins with the last common ancestor algorithm of Diamond after protein prediction with Augustus. Number of portin is shown for each taxa. The ratio of taxonomic rank of annotations across AMF lineages is shown in a pie chart.

## Re-use potential

Current microbiology investigations are focused on addressing the factors shaping the structure of microbial communities. To drive the development of tomorrow’s biotechnology it is essential to understand biological pathways both at the organism level and at the inter-microbial relationships scale, for prokaryotic and eukaryotic organisms together. This dataset provides a more complete and comprehensive view of the pool of genes and proteins, genetic diversity and distribution of eukaryotic microbes in soil and plant-associated microbiomes. At the molecular level, the use of this data is relevant to address biological questions in both fundamental research on plant-microbe interactions and applied, agronomical research, such as the study of potential metabolic functions of telluric eukaryotes, or of the interaction pathways between microbial members of the community. Moreover, the more accurate taxonomic annotation provides an unparalleled opportunity to assess how microbial eukaryotes are distributed across the soil and plant-associated microbial-environments. As illustrated in the data validation section, this improvement of microbial eukaryote representation has allowed us to increase by a factor of six times the detection of the ubiquitous AMF species, which are of high agronomic and economic interest.

Any research involving study of soil eukaryotes from evolutionary research on gene flow and transfers within the biome to more translational research aiming at deciphering important soil functions and biochemical pathways will benefit from this improved dataset of soil proteins with up to date taxonomic annotation. In addition, our data can be cross-referenced with the metadata provided by the JGI (downloadable from the IMG/M portal) which includes geo-tracking and a wealth of environmental, sampling and processing information on each metagenome. They can be linked to proteins and annotations by searching for the ‘metagenomeID’, as each protein name in our dataset has a semantic structure based on the following pattern: ‘contigName_metagenomeID.geneID’, to offer this possibility. On one hand, the ecological metadata provides the unprecedented potential to study the effect of the environment on community structures and to have a better, more comprehensive view on how external factors influence the eukaryotic soil microbial communities. On the other hand, metadata on sampling and processing could be useful to assess which parameters affect the diversity and sequencing of eukaryotes in metagenomes and help to shape future protocols. Moreover, in this study, we provide a fully documented pipeline and protocol available as python scripts from the detection of putative eukaryotic contigs to the *ab initio* model selection for Augustus gene prediction and further Diamond-based taxonomic annotation, that can be re-used to improve the annotation of eukaryotes on any microbiome data, including in other biomes than the soil.

## Conclusion

We produced a high quality *de novo* protein prediction and accurate taxonomic assignment of soil eukaryotic microorganisms. We predicted ca. 8.3 million eukaryotic proteins, of which 5,615,822 were present on 3.1 million non-chimeric high confidence eukaryotic contigs by mining 6,873 soil and plant-associated assembled metagenomes from the JGI’s IMG/M server using Augustus, a eukaryotic protein predictor. Comparing this protein prediction with the original JGI one, we showed that our pipeline provides a more complete and accurate representation of soil eukaryotic proteins. This new set of proteins better represents the eukaryotic genetic content from soil and plant-associated metagenomes and has been assigned a more reliable taxonomic annotation. Consistent with previous studies, most eukaryotic proteins in the soil are of fungal origin, but several protist taxa are also represented. Hence this dataset represents a unique new resource of interest in studying soil microbial ecosystems, both at the structure and function level.

## Availability of source code and requirements (optional, if code is present)

- Project name: EukaProt_in_PublicSoilMetag
- Project home page: https://github.com/CaroleBelliardo/EukaProt_in_PublicSoilMetag.git
- Operating system(s): Platform independent
- Programming language: Python3
- Other requirements: Python3.8 or higher
- License: License: GNU General Public License v3.0

## Availability of supporting data and materials

Fasta files are availaible in [’Fasta’](https://data.inrae.fr/privateurl.xhtml?token=5fe3ff97-c716-41ff-98a0-f6e9aa0631d3) repository, and taxonomic annotations in [’Taxonomy’](https://data.inrae.fr/privateurl.xhtml?token=5fe3ff97-c716-41ff-98a0-f6e9aa0631d3) repository.

## Abbreviations

AMF: Arbuscular mycorrhizal fungi
bp: base pairs
Kb: Kilobases
Mb: Megabases
MAG: Metagenome-assembled genomes
IMG/M: the Integrated Microbial Genomes & Microbes
JGI: Joint Genome Institute
TaxID: NCBI Taxonomy Identifier

## Declarations

### Consent for publication

“Not applicable”

### Funding

This project was supported by the Mycophyto company and the INRAE plant’s Health department.

## Acknowledgements

We thank all members of the bioinformatic platform of the Institute Sophia Agrobiotech, Sophia Antipolis, France, for they help and support. We also want to warmly thank Samuel Mondy PhD., INRAE, Dijon, France for his advices on soil microbial analyses.

## Notes

### Competing Interest Statement

The authors have declared no competing interest.

### Summary of Updates

Corrected affiliation of authors

